# Connectome-based predictive modeling of individual anxiety

**DOI:** 10.1101/2020.01.30.926980

**Authors:** Zhihao Wang, Katharina S. Goerlich, Hui Ai, André Aleman, Yuejia Luo, Pengfei Xu

## Abstract

Anxiety-related illnesses are highly prevalent in human society. Being able to identify neurobiological markers signaling high trait anxiety could aid the assessment of individuals with high risk for mental illness. Here, we applied connectome-based predictive modeling (CPM) to whole-brain resting-state functional connectivity (rsFC) data to predict the degree of anxiety in 76 healthy participants. Using a computational “lesion” method in CPM, we then examined the weights of the identified main brain areas as well as their connectivity. Results showed that the CPM could predict individual anxiety from whole-brain rsFC, especially from limbic areas-whole brain and prefrontal cortex-whole brain. The prediction power of the model significantly decreased from (simulated) lesions of limbic areas, lesions of the connectivity within the limbic system, and lesions of the connectivity between limbic regions and the prefrontal cortex.

Although the same model also predicted depression, anxiety-specific networks could be identified independently, centered at the prefrontal cortex. These findings highlight the important role of the limbic system and the prefrontal cortex in the prediction of anxiety. Our work provides evidence for the usefulness of connectome-based modeling of rsFC in predicting individual personality differences and indicates its potential for identifying personality structures at risk of developing psychopathology.

## Introduction

Anxiety, a personality dimension in healthy individuals, is a universal negative emotion that entails avoidance behaviors such as worrying, irritability, difficulty to relax and a tendency to interpret ambiguous situations as threatening (Spielberger, 1983; Grachev and Apkarian, 2000; Eysenck et al., 2007). Although anxiety has valuable adaptive benefits, individuals enduring high trait anxiety may be at risk to develop mental disorders (Pezawas et al., 2005; Sandi and Richter-Levin, 2009; Cremers et al., 2010). It has been estimated that an alarming 28.8% of the general population suffer from an anxiety-related disorder at some point in their lifetime (Kessler et al., 2005). Identifying neurobiological markers signaling high trait anxiety could aid the assessment of high-risk individuals, especially those with difficulty expressing their feelings to others (Eisenberger et al., 2005; Drysdale et al., 2017).

Anxiety is a complex emotion that is associated with mutual inhibition between subcortical and cortical areas of the brain (Tovote et al., 2015; Xu et al., 2019). Particularly, the limbic system and the prefrontal cortex are highly engaged in the balance of these mutually inhibitory relationships (Hofmann et al., 2012). The limbic system has long been considered crucial for emotion processing (Fuchs and Flügge, 2003), and the prefrontal cortex plays an important role in top-down regulation of limbic activity (Mochcovitch et al., 2014). Regarding anxiety disorders, an emotion dysregulation model has been proposed that is characterized by an imbalance of prefrontal regulation over limbic activity (Behar et al., 2009; Manber Ball et al., 2013; Mochcovitch et al., 2014). Specifically, it has been suggested that patients with anxiety disorders experience chronic hyper-arousal in the limbic circuitry (Kim et al., 2011). This fatigues the top-down regulation system in the prefrontal cortex, resulting in ineffective control particularly of negative emotions (Manber Ball et al., 2013). With regard to trait anxiety as a personality dimension in non-clinical samples, it has been demonstrated that anxiety-prone individuals require greater engagement of prefrontal regions to down-regulate negative emotions (Campbell-Sills et al., 2011). In spite of this, no effort has been made as yet to predict anxiety at the individual level for healthy people. Individualized prediction of anxiety would advance our understanding of the underlying neural mechanism and facilitate early identification of proneness to clinical anxiety.

Recently, a novel approach has been tested to account for inter-individual variability in brain functional networks: Connectome-based predictive modeling (CPM). CPM is a novel data-driven approach for developing predictive models of brain–behavior relationships that can detect individual variability more accurately (“functional connectome fingerprinting”; Finn et al., 2015) by extracting and summarizing the most relevant features from resting-state functional connectivity (rsFC) using full cross-validation (Shen et al., 2017). Despite testing a specific hypothesis, CPM provides more holistic measures with whole-brain analyzes. This approach has successfully been used to predict individual personality traits and aspects of functional cognition, such as fluid intelligence (Finn et al., 2015), sustained attention (Rosenberg et al., 2015), and creative ability (Beaty et al., 2018). Moreover, a computational lesion method based on CPM can reveal brain regions that are important in individualized prediction (Feng et al., 2018). The computational lesion method allows for the computational manipulation of brain regions to be “lesioned”, and thus represents a non-invasive method simulating the effect of lesions in particular brain regions on aspects of neuropsychological functioning.

The aim of current study was to predict individual levels of anxiety in healthy participants by applying CPM and the computational lesion approach to whole-brain resting-state fMRI data. We hypothesized that CPM would successfully predict anxiety at the individual level. Based on the emotion dysregulation model (Behar et al., 2009) and the computational lesion method in CPM (Feng et al., 2018), we hypothesized that both the limbic system and the prefrontal cortex would significantly contribute to the identification of individual levels of anxiety.

## Materials and Methods

### Participants

Eighty-eight healthy undergraduate students took part in MRI scanning. All the participants had no history of neurological and psychiatric disorders or head injury. After excluding excessive head motion [10 participants, exceeding 2.5 mm maximum translation, 2.5° rotation or 0.2mm mean frame-wise displacement (FD; Yan et al., 2013; Power et al., 2014)] and outliers in Back Anxiety Inventory scores (BAI: 2 participants, out of mean ± 2.5 std), the final sample consisted of 76 participants (38 females; age = 21.34 ± 1.76). The study was approved by the local Ethics Committee at Beijing Normal University and written informed consent was obtained from all participants.

### Anxiety assessment

To assess anxiety, we used the Chinese version of the BAI (Aaron T. Beck et al., 1988). This inventory consists of 21 items, each answer being scored on a four-point Likert scale of 1 (not at all) to 4 (severely). Participants also completed the Chinese version of the Zung self-rating depression scale (SDS; Zung, 1965).

### Image acquisition

MRI data were acquired with a Siemens Trio 3T scanner powered with Total Imaging Matrix technique at the Imaging Center for Brain Research at Beijing Normal University. Both the fMRI and high-resolution 3D structural brain data were obtained using a 12-channel phased-array head coil with implementing a parallel imaging scheme that generalized auto-calibrating partially parallel acquisitions (Griswold et al., 2002). The fMRI data were acquired with a gradient-echo echo-planer imaging sequence with the following parameters: repetition time (TR) = 2000 ms, echo time (TE) = 30 ms, 33 transversal slices, slice thickness 3.5 mm with gap 0.7 mm, flip angle = 90°, field of view (FOV) = 224 mm × 224 mm, data matrix = 64 × 64, 240 volumes scanned in 8 min, and spatial coverage (3.5 + 0.7) mm/slice×33 slices ≈ 139 mm. Additionally, the 3D structural brain images (1mm^3^ isotropic) were acquired for each participant using a T1-weighted 3D magnetization-prepared rapid gradient echo sequence with the following parameters: TR / TE = 1900 ms / 3.44 ms, flip angle = 9°, data matrix = 256 × 256, FOV = 256 mm × 256 mm, BW = 190 Hz / pixel, 176 image slices along the sagittal orientation, obtained in about 6 min. During resting-state scanning, all participants were instructed just to keep still, close their eyes, remain awake and think of nothing in particular.

### Preprocessing

Functional MRI data were preprocessed with DPABI (http://rfmri.org/dpabi; (Yan et al., 2016), a software package based on SPM12 (version no.7219; https://www.fil.ion.ucl.ac.uk/spm/software/spm12/). It comprised the following steps: 1) discarding the first 10 volumes to decrease the signal’s instability; 2) correcting slice timing; 3) realignment; 4) co-registering the T1-weighted image to the corresponding mean functional image; 5) segmenting into grey matter, white matter and cerebrospinal fluid by DARTEL; 6) regressing common nuisance out by Compcor (Behzadi et al., 2007), including the white matter signal, the cerebrospinal fluid signal, 24 movement regressors, and global signal. The 24 movement regressors included autoregressive models of motion incorporating six head motion parameters, six head motion parameters one time point before, and the 12 corresponding squared items (Friston et al., 1996; Yan et al., 2013). Note that we administrated global signal regressors to eliminate noise, including head motion and respiratory as well as cardiorespiratory artifacts (Power et al., 2015; Ciric et al., 2017, 2018; Murphy and Fox, 2017). 7) detrending; 8) normalizing to the standard Montreal Neurological Institute space (MNI template, resampling voxel size 3×3×3 mm^3^); 9) smoothing with a Gaussian kernel of 4 mm full width at half maximum (FWHM); 10) filtering (0.01-0.1Hz).

### Functional Network Construction

After preprocessing, we constructed the static whole-brain rsFC matrices. Specifically, we employed a 268-node functional atlas that was derived from a group-wise spectral clustering algorithm. This group-wise optimization approach ensures functional homogeneity within each subunit, and time-course consistency of the network nodes at the group level, resulting in improved reliability and sensitivity of the network analyses (Shen et al., 2010, 2013). In light of previous literatures using this atlas (Shen *et al*., 2013; Feng *et al*., 2018), nodes were further divided into ten lobes, including prefrontal lobe (46 nodes), motor lobe (21 nodes), insula lobe (7 nodes), parietal lobe (27 nodes), temporal lobe (39 nodes), occipital lobe (25 nodes), limbic lobe (36 nodes), cerebellum lobe (41 nodes), subcortical lobe (17 nodes) and brainstem lobe (9 nodes; Fig. 2d). After Fisher’s Z transformation, the resulting 268 ×268 symmetric matrices represented edges from the rsFC profile.

### Connectome-based predictive modeling (CPM)

Based on the whole-brain rsFC, we used CPM to predict the degree of anxiety individually. In light of ten simple rules for applying predictive modeling to rsFC data (sample size < 200; Scheinost *et al*., 2019) and to be consistent with past work employing CPM (Finn et al., 2015; Rosenberg et al., 2015; Beaty et al., 2018), we performed leave-one-out cross-validation (LOOCV). That is, in each iterative analysis, the predictive model was built based on n - 1 participants (training set) and then the score of the remaining participant (test set) was predicted. In this study, we ran it 76 times. Steps were as follows (Fig. 1). 1) Data preparation, preparing rsFC matrices and BAI scores of all participants. 2) Edge selection, performing Pearson correlations between each edge in rsFC matrices and BAI scores in the training set and selecting only the most significantly correlated edges as predictive networks. The threshold was set at *p* < 0.01 (Dosenbach et al., 2010; Rosenberg et al., 2015; Takagi et al., 2018). 3) Network construction, a positive network (i.e., positive correlation within selected edges) and a negative network (i.e., negative correlation within selected edges). 4) Network strengths, summing the edges of the positive network and the negative network separately for each participant. 5) Model building, fitting the linear model between BAI scores and network strengths across the training set for both the positive network and the negative network to build the positive model and the negative model, respectively. 6) Individual prediction, applying models to predict the BAI score according to the network strengths in the test set (the excluded participant). Notably, normalization of each edge across the training set was performed and parameters from the training set were applied to the test set.

**Fig.1.**
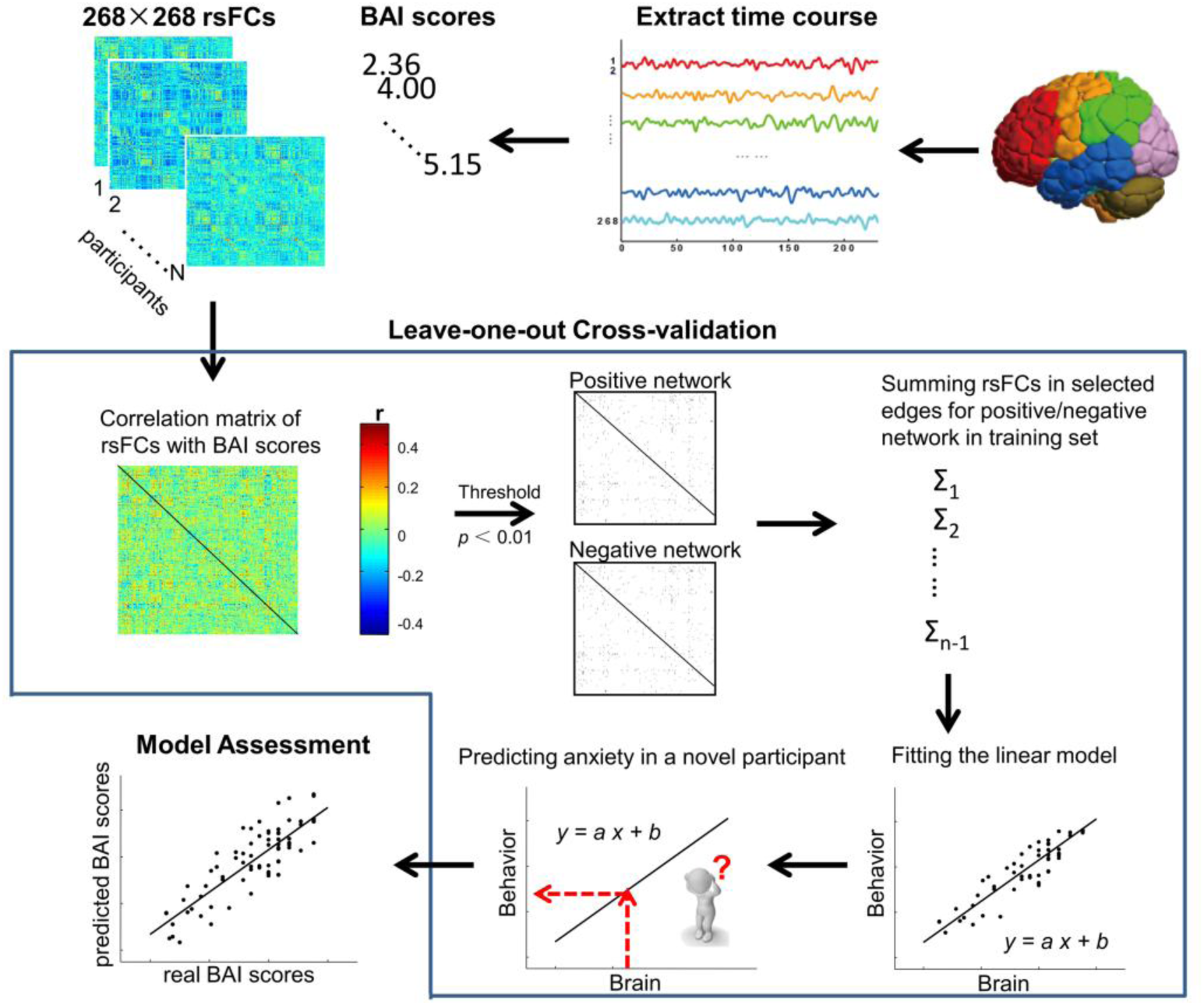
The schematic flow of connectome-based predictive modeling. Note: rsFCs, resting-state functional connectivity; BAI, Beck Anxiety Inventory.

### Model assessments

After LOOCV, the model validity was evaluated by Pearson correlation coefficient (r) between real BAI scores and predicted BAI scores for the positive and negative model, respectively. To check whether these effects were expected by chance, we further ran permutation test for 1000 times. Specifically, permutation tests randomly shuffled the label between real BAI scores and rsFC matrices each time and then calculated a correlation coefficient from CPM. The calculating 1000 correlation coefficients composed null distributions. We then calculated the percentile of model validity value among these null distributions.

### Computational lesion prediction

Because the obtained networks were slightly different in each iteration, we extracted the common edges (i.e., overlaps of all the negative networks) to construct common networks and then categorized them into 10 macroscale lobes. After that, lesion prediction analyses were performed to examine the weights of the limbic lobe and the prefrontal lobe as well as their connectivity. For example, after excluding edges in the limbic lobe (36 nodes), the remaining 232 × 232 rsFC matrices were used to predict individual anxiety (Feng *et al*., 2018). Finally, we contrasted whole-brain prediction with lesion prediction in terms of predictive power by Steiger’s Z (Steiger, 1980).

### Control analyses

To avoid potential confounding effects of head movements, gender and age, control analyses were conducted. We repeated our analyses by implementing scrubbing with the criterion of an FD above 0.2 mm (Jenkinson et al., 2002; Yan et al., 2013). Additionally, partial Pearson correlation analyses were conducted with mean FD of head motion (Jenkinson et al., 2002), gender and age as co-variables in edge selection and final correlation, separately.

### Distinct brain networks between anxiety and depression

Given that anxiety was highly correlated with depression (Stavrakaki and Vargo, 1986; Clark and Watson, 1991), we explored distinct brain networks for anxiety versus depression in this module (Fig. 5). After excluding two participants (out of mean ± 2.5std) in BAI scores, 46 high anxious participants were selected from the criteria of the top 55% in BAI scores, and 44 high depressive participants were selected from the top 55% in SDS scores (overlap: 35 participants). Eleven anxiety-specific participants (BAI: 25∼32, mean ± std = 26.20 ± 2.15; SDS: 20∼31, mean ± std = 28.00 ± 3.23) and nine depression-specific participants (BAI: 21∼24, mean ± std = 23.11 ± 1.05; SDS: 32∼38, mean ± std = 34.00 ± 1.87) were identified for further explorative contrast analyses after excluding 35 anxiety participants comorbid with depression.

We employed a CPM-like model to explore distinct brain networks between anxiety and depression (Takagi *et al*., 2018; Fig. 5). In an LOOCV manner, we conducted two-sample *t*-tests between the anxiety-specific and the depression-specific rsFC matrices across the training set for each edge. Based on the significance threshold *p* < 0.01 (Dosenbach *et al*., 2010; Rosenberg *et al*., 2015; Takagi *et al*., 2018), anxiety-specific edges were selected to construct the anxiety-specific network across the training set for those anxiety-specific edges significantly higher than depression-specific edges. Likewise, depression-specific edges were selected to construct the depression-specific network across the training set for those depression-specific edges significantly higher than anxiety-specific edges. Next, the “anxiety score” of the remaining participant (test set) was defined summing rsFCs in the anxiety-specific network; the “depression score” was calculated correspondingly.

Then, we validated the difference between anxiety-specific and depression-specific networks by performing a two-sample *t*-test between “anxiety scores” and “depression scores”. Lastly, we extracted common edges (i.e., overlaps of all anxiety-specific networks and overlaps of all depression-specific networks) to construct common networks and divided them into 10 macroscale lobes.

## Results

### Normality test

BAI scores were not normally distributed according to the Kolmogorov-Smirnov Test of Normality (BAI: Z = 1.695, *p*= 0.006, range: 21∼44, mean ± std = 27.78 ± 6.21; SDS: Z = 0.788, *p* = 0.564, range: 20∼57, mean ± std = 34.34 ± 7.89). Therefore, we performed reciprocal transformation (100 / scores; Box and Cox, 1964) to normalize BAI scores (BAI: Z = 1.312, *p* = 0.064; 3.75 ± 0.70).

### CPM assessments of anxiety

LOOCV revealed that the negative model, but not the positive model, had significant prediction power to anxiety scores [positive model, r _(74)_ = −0.01, *p* = 0.939, Fig. 2a; negative model, r _(74)_ = 0.33, *p* = 0.004, Fig. 2b]. Permutation tests (1000 times) revealed that this result was expected above chance (*p* = 0.036, Fig. 2c). Thus, subsequent analyses focused on the negative model. Please note that the negative model indicated positive correlations between rsFC and levels of anxiety because of reciprocal transformation.

**Fig. 2.**
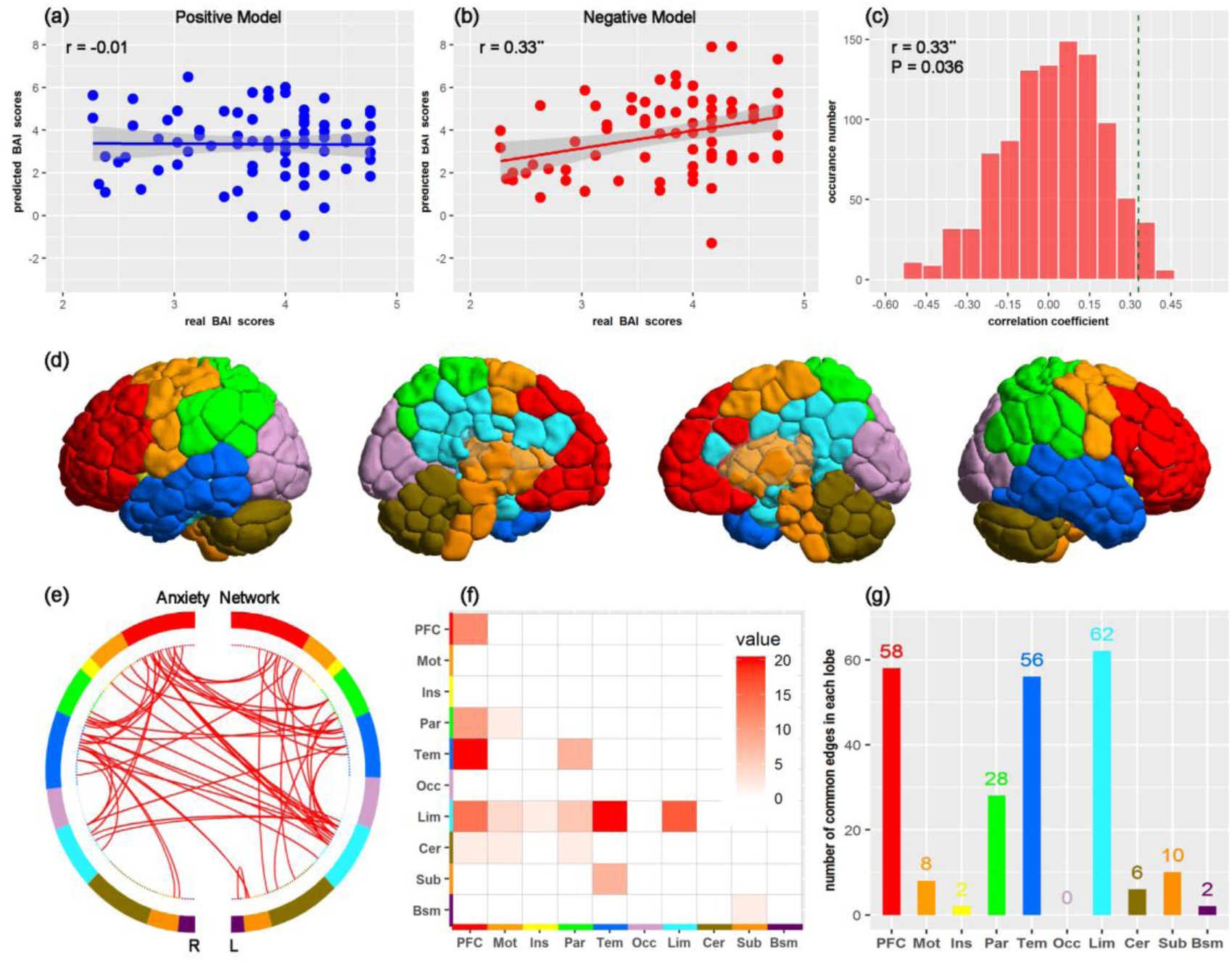
Correlations between real BAI score and predicted BAI scores in (a) positive model and (b) negative model. (c) Permutation distribution of correlations in the negative model. (d) Whole brain parcellation with 10 lobes. (e) Common edges contributing to anxiety prediction. (f) (g) Division of anxiety network into 10 lobes and the contribution of each lobe to anxiety prediction. Note: BAI, Beck Anxiety Inventory; PFC, prefrontal; Mot, motor; Ins, insula; Par, parietal; Tem, temporal; Occ, occipital; Lim, limbic; Cer, cerebellum; Sub, subcortical; Bsm, brainstem. R: right; L, left. \*\**p*<0.01.

The number of edges, common edges and networks across LOOCV are shown in Table 1 and Fig. 2e. Importantly, prediction through the common networks resulted in a stronger correlation than the whole-brain network prediction (r _(74)_ = 0.85, *p* = 0; Steiger’s Z = 8.09, *p* < 0.001). It suggested that the common edges have a stronger prediction power despite the small number of common edges. The number of edges in each lobe from common networks is shown in Fig. 2f & g. Furthermore, the top 10 highly connected brain nodes were located in the ventral anterior cingulate cortex (vACC), ventral-lateral prefrontal cortex (vlPFC), posterior cingulate cortex (PCC), medial superior parietal gyrus (mSPG), inferior temporal gyrus (ITG) and middle temporal gyrus (MTG; Fig. 3).

**Fig. 3.**
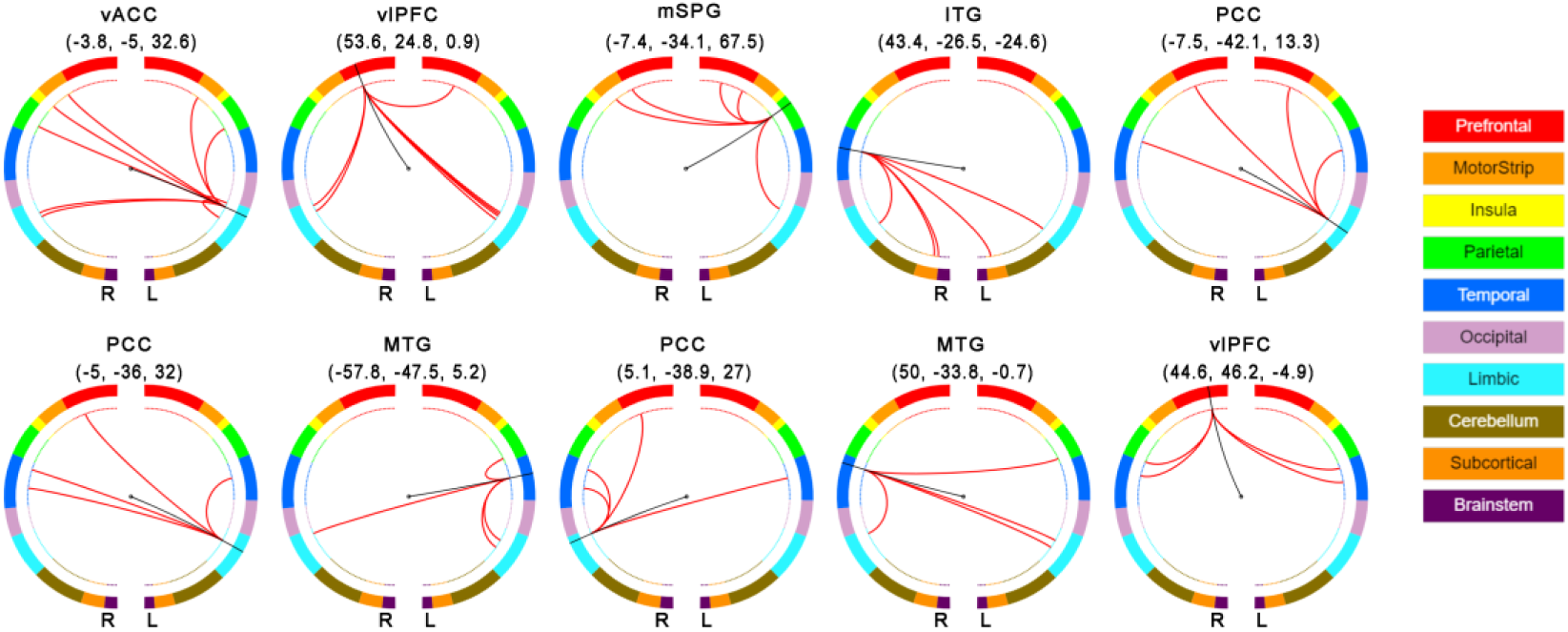
Connectivity patterns of the top 10 most highly connected brain nodes. The MNI coordinates are shown under the name of each brain node. Note: vACC, ventral anterior cingulate cortex; vlPFC, ventral-lateral prefrontal cortex; mSPG, medial superior parietal gyrus; ITG, inferior temporal gyrus; PCC, posterior cingulate cortex; MTG, middle temporal gyrus; R, right; L, left.

**Table 1.** The number of edges or networks across LOOCV in CPM of anxiety, anxiety-specific networks and depression-specific networks

|  | CPM of anxiety | anxiety-specific<br>networks | depression-specific<br>networks |
| --- | --- | --- | --- |
| edges [min, max] | [130, 169] | [108, 197] | [93, 179] |
| common edges | 58 | 54 | 40 |
| networks | 76 | 20 | 20 |

Note: CPM, connectome-based predictive modeling.

### Computational lesion prediction

In addition to the limbic lobe and the prefrontal lobe, we also conducted lesion prediction analyses for the temporal lobe, because the number of common edges in the temporal lobe was almost as high as for the limbic lobe and the prefrontal lobe (Fig. 2g). These results showed that the anxiety prediction power of the model decreased dramatically for lesions to the limbic or temporal lobes, for lesioned connectivity within the limbic lobe, and for lesioned connectivity between the limbic and the prefrontal lobe (see Table 2 and Fig. 4).

**Table 2.** Lesion predictions

| Lesion | Predictive power |  | Difference from the whole-brain model<br>Steiger's Z |
| --- | --- | --- | --- |
|  | r | p |  |
| Lim | 0.16 | 0.165 | -2.38* |
| PFC | 0.24 | 0.037 | -1.44 |
| Tem | 0.17 | 0.141 | -2.24* |
| Lim-Lim | 0.29 | 0.012 | -3.06** |
| Lim-PFC | 0.28 | 0.014 | -2.03* |
| Lim-Tem | 0.29 | 0.012 | -1.21 |
| PFC-PFC | 0.33 | 0.004 | 0 |
| PFC-Tem | 0.32 | 0.005 | -0.36 |
| Tem-Tem | 0.34 | 0.002 | 1.87 |

Note: Lim, limbic; PFC, prefrontal; Tem, temporal; \* $p < 0.05$ ; \*\* $p < 0.01$ .

**Fig. 4.**
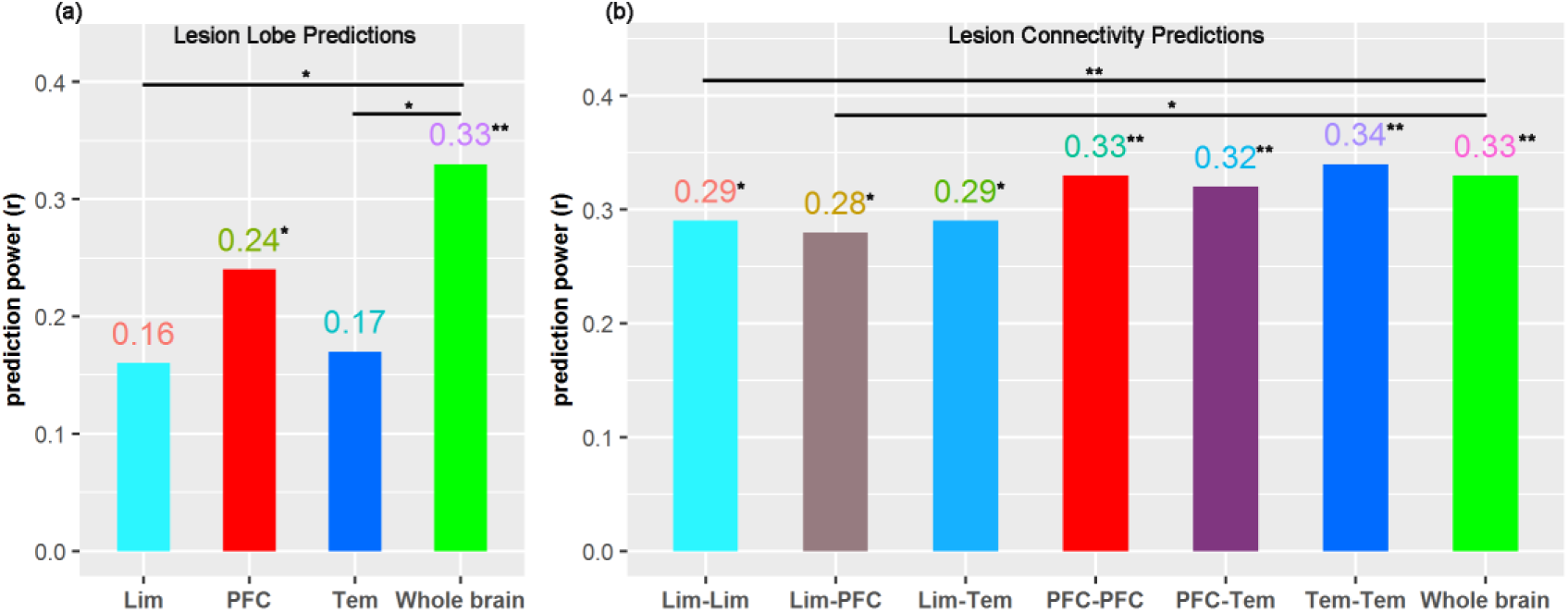
Lesion predictions. (a) Lesion lobe predictions. (b) Lesion connectivity predictions. Note: Lim, limbic; PFC, prefrontal; Tem, temporal. \**p*<0.05, \*\**p*<0.01.

**Fig. 5.**
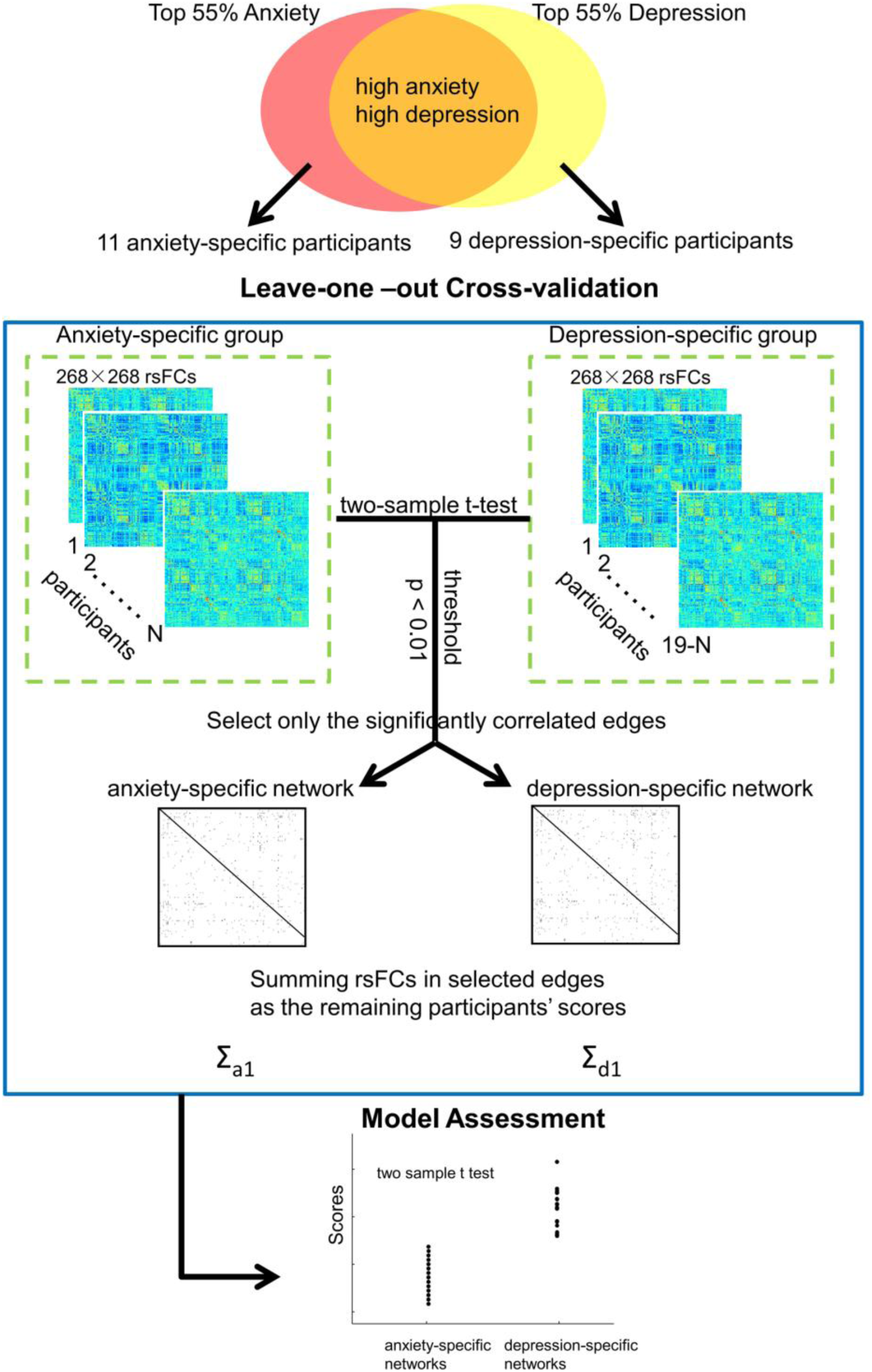
The schematic flow for exploring distinct brain networks between anxiety and depression. Note: rsFCs, resting-state functional connectivity.

### Control analyses

After controlling for nuisance variables in edge selection and final correlation, and for scrubbing in preprocessing, the predictive models remained significant (Table 3). The common networks drawing from anxiety could also predict individual depression through CPM [r _(74)_ = 0.64, *p* < 0.001, Fig. 6b].

**Table 3.** Control analyses with adding head motion, age and gender as covariates in edge selection and final correlation, as well as scrubbing of head motion.

|  | Control at edge selection | Control at final correlation |
| --- | --- | --- |
| Head motion | $r = 0.31, p = 0.006$ | $r = 0.33, p = 0.004$ |
| Age | $r = 0.24, p = 0.035$ | $r = 0.30, p = 0.008$ |
| Gender | $r = 0.22, p = 0.051$ | $r = 0.32, p = 0.005$ |
| Scrubbing | $r = 0.28, p = 0.015$ | |

**Fig. 6.**
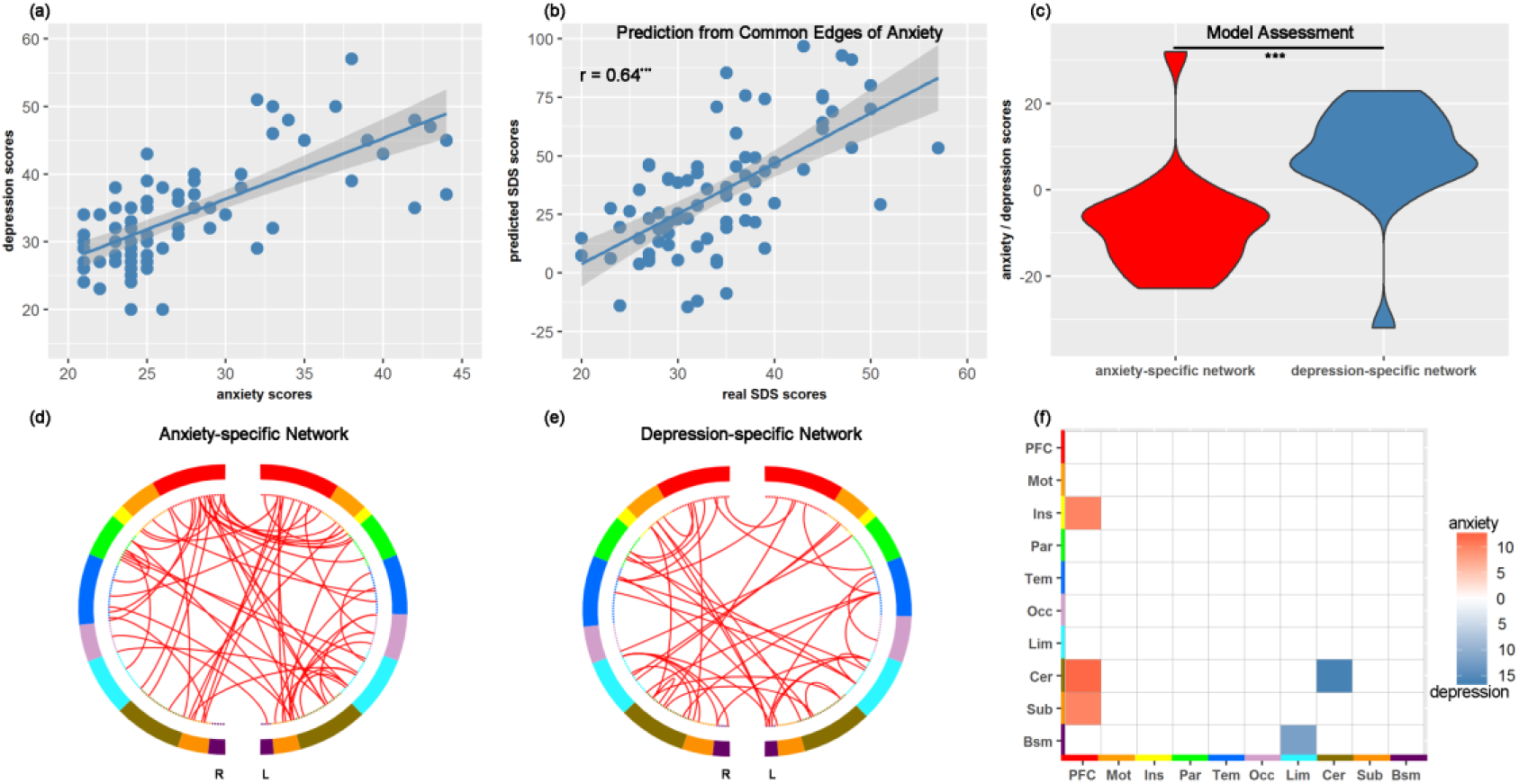
(a) Distribution of anxiety scores and depression scores for each participant. (b) Predicting depression by common edges obtained from anxiety. (c) Anxiety scores obtained from anxiety-specific networks, and depression scores obtained from depression-specific networks for two sample *t*-test analysis to validate these networks. For the purpose of visualization, anxiety scores and depression scores were normalized. (d) anxiety-specific network. (e) depression-specific network. (f) Division of anxiety-specific network and depression-specific network into 10 lobes. Note: PFC, prefrontal; Mot, motor; Ins, insula; Par, parietal; Tem, temporal; Occ, occipital; Lim, limbic; Cer, cerebellum; Sub, subcortical; Bsm, brainstem. R: right; L: left. \*\*\**p*<0.001.

### Distinct brain networks between anxiety and depression

Given the strong correlation between (reciprocally transformed) BAI scores and SDS scores [r _(74)_ = −0.73, *p* < 0.001, Fig.6a], we explored distinct brain networks between anxiety and depression. The two-sample *t*-test revealed a significant difference between “anxiety scores” and “depression scores” [*t* _(38)_ = −4.15, *p* <0.001, Cohen’s *d* = −1.857, Fig. 6c], validating significant differences between the anxiety-specific and the depression-specific networks. The number of edges, common edges and networks across LOOCV for anxiety-specific and depression-specific networks are shown in Table 1, Fig. 6d and Fig. 6e. For the purpose of visualization, common networks in each lobe greater than or equal to ten were presented (Fig. 6f).

## Discussion

In this study, we employed CPM, a data-driven, full cross-validation approach, to predict levels of anxiety in healthy participants from whole-brain rsFC. We demonstrated that anxiety could be predicted by individual functional connectivity patterns, especially those from the limbic system and prefrontal cortex. More specifically, “lesion prediction” revealed that connectivity within the limbic system as well as connectivity between the limbic system and prefrontal cortex most significantly predicted individual anxiety levels. The nodes that highly contributed to the predictive model included ventral anterior cingulate cortex (vACC), ventral-lateral prefrontal cortex (vlPFC), posterior cingulate cortex (PCC), medial superior parietal gyrus (mSPG), inferior temporal gyrus (ITG) and middle temporal gyrus (MTG). We also explored anxiety-specific networks and depression-specific networks.

Using a computational lesion method, our CPM-based approach suggested that the anxiety-predictive power of the model significantly decreased with a) lesion of the limbic system, b) lesion of the connectivity within the limbic system, and c) lesion of the connectivity between the limbic system and prefrontal cortex (Fig. 4). These results, except increased prefrontal-limbic connectivity, conform to the emotion dysregulation model (Behar et al., 2009). It is common that anxiety is associated with emotional hyper-arousal, which was accompanied by hyper-connectivity within the limbic system (Kim et al., 2011). However, interpretations of the observed prefrontal-limbic hyper-connectivity are less straightforward. In anxiety-related disorders, attenuated prefrontal responses (Manber Ball et al., 2013) to over-responsiveness of the limbic system (Etkin et al., 2004; Qin et al., 2014; Bishop, 2007) have been observed. A nonclinical study found that anxiety-prone individuals were as successful as controls to reduce negative their emotions, but required greater engagement of lateral and medial PFC for successful down-regulation of negative emotions (Campbell-Sills et al., 2011). The increased prefrontal-limbic connectivity at rest in the current study might thus reflect more effort to regulate emotions in daily life in participants with higher degrees of anxiety.

Among the prefrontal-limbic circuit, vACC, vlPFC and PCC were key nodes that contributed to the anxiety-predictive model. vACC plays a key role in the modulation of physiological arousal (Maresh et al., 2013) and the control of negative affect (LeDoux, 2003; Passamonti *et al*., 2008; Petrovic *et al*., 2005). vlPFC has been implicated in effortful down-regulation of negative emotion (Campbell-Sills et al., 2011). The PCC is involved in mediating interactions of emotional and memory-related processes (e.g., Maddock *et al*., 2003). Taken together, these key nodes in the current study might reflect the neural underpinning of anxious individuals within the normal range experiencing a successful emotion regulation. However, we did not find the amygdala in the top ten connected brain nodes. It has long been acknowledged that the amygdala exerts a significant role in emotion regulation (Whalen et al., 2002; Somerville et al., 2004; Sylvester et al., 2012; Grupe and Nitschke, 2013; Tovote et al., 2015; Xu et al., 2019). Despite this, previous hypothesis-driven studies may over-represent the amygdala. Specifically, the significant diagnostic effect of amygdala in ROI studies disappeared when whole-brain studies were also considered in a meta-analysis for mental disorders (including anxiety disorders; Sprooten *et al*., 2017). Another reason for failure to find the amygdala might be that it was difficult to infer the specific involvement of amygdala in the 268-node functional atlas, because the averaged size of the amygdala (1.24 cm^3^) is smaller than the averaged size of the other nodes (4.8 cm^3^; Brabec *et al*., 2010; Hsu *et al*., 2018). Future studies could attempt to account for such differences in the average size of relevant brain regions.

Surprisingly, anxiety-prediction power also significantly decreased with lesion of the temporal gyrus. Within the temporal gyrus, medial (MTG) and inferior temporal gyrus (ITG) were highly connected brain nodes in anxiety-predictive networks. MTG plays a crucial role in social perception to uncertain threats (Haxby et al., 2002; Geng et al., 2018; Feng et al., 2019). ITG is associated with the ventral visual pathway (Baddeley et al., 1997). Its volume is associated with anxiety disorders, functionally reflecting the processing of external social cues (Liao et al., 2011). Therefore, the current results might reflect the information flow of social perception to uncertain threats from ITG to MTG in anxiety.

It is important to note that anxiety and depression were highly correlated in this study (a common finding) and that the anxiety-predictive model also predicted depression. Anxiety in this study may therefore be considered as referring to an anxious-depressed trait. Such close connection is not surprising given the high correlation of anxiety and depression in real life (Stavrakaki and Vargo, 1986; Clark and Watson, 1991). It has been demonstrated that anxiety-prone individuals require more engagement of prefrontal cortex to accomplish down-regulation of negative emotion (Campbell-Sills et al., 2011), which has also been found in depressed individuals (Johnstone et al., 2007). The neuropsychological mechanisms during emotion regulation may thus be similar in anxiety and depression, especially regarding the prefrontal-limbic connectivity (Campbell-Sills et al., 2011).

We also preliminarily dissociated anxiety and depression at the network level. Anxiety-specific networks were mainly centered in the prefrontal cortex, connecting with the insula, subcortical lobes and cerebellum, whereas depression-specific networks were mainly observed within the cerebellum and between the limbic system and brainstem (Fig. 6f). However, the sample size of the high anxiety-specific and high depression-specific participants was too small in the current study; these results should thus be interpreted with caution. It has been documented that differentiation of anxiety and depression is difficult because of their high inter-correlation (Stavrakaki and Vargo, 1986; Clark and Watson, 1991). Accordingly, categorical definitions of anxiety and depression are difficult to capture the distinctions of brain abnormalities (Oathes et al., 2015; Pannekoek et al., 2015). Nonetheless, the current study applied a novel brain model to understand the neurobiological distinction of anxiety and depression. Future studies with large sample sizes of each specific group are needed.

The present work represents advances in neuroscience to advocate the applications of rsFC in a cross-validation manner (Shen et al., 2017). An individualized predictive model of anxiety was estimated from whole-brain rsFC profiles. The advantages of this approach are fourfold. First, resting-state data is easy to collect relative to task-dependent experiments (Rosazza and Minati, 2011). Despite being task-free, rsFC can track individual cognitive performance (Scheinost et al., 2019), even higher-order functions such as trust (Lu et al., 2019). Importantly, rsFC is stable (even across three years; Horien et al., 2019) and unique (functional connectome ‘fingerprinting’; Finn et al., 2015). Second, compared to standard group comparisons between high-anxious and low-anxious individuals, the present model is more sensitive for the assessment of aspects of neuropsychological functioning in the individual brain (Gabrieli et al., 2015; Dubois and Adolphs, 2016). Although studies with correlation or regression models also used the term “prediction”, such in-sample prediction rather than out-of-sample prediction, incurs over-fitting (Gabrieli et al., 2015; Dubois and Adolphs, 2016). Third, instead of region-of-interest (ROI) or seed-based rsFC studies, we were able to obtain an objective picture of whole-brain rsFC networks in individual participants, eliminating the risk of over-representation of certain ROIs (Sprooten et al., 2017). In this way, our findings were still meaningful, conforming to emotion regulation network regarding ‘emotion dysregulation model’ (Behar et al., 2009; Manber Ball et al., 2013; Mochcovitch et al., 2014). Finally, the approach of implementing computational (‘virtual’) lesions may have advantages above noninvasive neurostimulation methods, for instance, tDCS and TMS, which can cause some discomfort to participants. And although the causal explanatory power of neurostimulation techniques is stronger, they have significant drawbacks, the most important one being that they can only target superficial cortical regions as the depth of stimulation is very limited. A combination of the strenghts of both methods can also be indicative. Indeed, the computational lesion approach might provide potential neurostimulation targets (i.e., network-guided TMS; Fox *et al*., 2012) for future studies to reduce the tension of anxiety.

Several limitations of the present study are worth mentioning. First, we used a group-level atlas in the individualized prediction of anxiety, which might overlook subtle brain-behavior associations if they are highly variable. It has been documented that individualized prediction with the group-level atlas has a weaker predictive power than that with the individualized template (Wang et al., 2018). Second, for the purpose of early identification in subclinical population, we only recruited healthy participants. Given the asymmetric mechanisms between patients and healthy participants (Liao et al., 2010), however, whether this result could extend to clinical diagnosis in anxiety disorders remains unknown. Future studies could test our model in patients with anxiety disorders.

To conclude, we established a brain connectivity-based model that was able to predict anxiety in novel individuals. In light of emotion dysregulation models, we demonstrated that regions of emotion regulation networks, such as vACC, vlPFC and PCC, were of importance for individual anxiety-prediction in healthy people. We also provide preliminary evidence for distinct networks in the classification of anxiety and depression. The current work may have important implications for the early identification of individuals enduring high trait anxiety in non-clinical populations.

## Acknowledgements

This work was supported by the National Natural Science Foundation of China (Key Project for International Collaboration 3192010300, 31530031, 31871137, 31700959 and 31671133, 31500920), Young Elite Scientists Sponsorship Program by China Association for Science and Technology (2018), Guangdong Key Basic Research Grant (2018B030332001), Guangdong young Innovative Talent Project (2016KQNCX149), Guangdong Pearl River Talents Plan Innovative and Entrepreneurial Team grant (2016ZT06S220), Shenzhen Science and Technology Research Funding Program (JCYJ20180507183500566 and CYJ20170412164413575) and Shenzhen Peacock Program (827-000235, KQTD2015033016104926).

